# Xolik: finding cross-linked peptides with maximum paired scores in linear time

**DOI:** 10.1101/155069

**Authors:** Jiaan Dai, Wei Jiang, Fengchao Yu, Weichuan Yu

**Affiliations:** Department of Electronic and Computer Engineering, The Hong Kong University of Science and Technology, Hong Kong, China; Division of Biomedical Engineering, The Hong Kong University of Science and Technology, Hong Kong, China.

## Abstract

**Motivation:** Cross-linking technique coupled with mass spectrometry (MS) is widely used in the analysis of protein structures and protein-protein interactions. In order to identify cross-linked peptides from MS data, we need to consider all pairwise combinations of peptides, which is computationally prohibitive when the sequence database is large. To alleviate this problem, some heuristic screening strategies are used to reduce the number of peptide pairs during the identification. However, heuristic screening criteria may ignore true findings.

**Results:** We directly tackle the combination challenge without using any screening strategies. With the additive scoring function and the data structure of double-ended queue, the proposed algorithm reduces the quadratic time complexity of exhaustive searching down to the linear time complexity. We implement the algorithm in a tool named Xolik, and the running time of Xolik is validated using databases with different number of proteins. Experiments using synthetic and empirical datasets show that Xolik outperforms existing tools in terms of running time and statistical power.

**Availability:** Source code and binaries of Xolik are freely available at http://bioinformatics.ust.hk/Xolik.html.

**Supplementary information:** Supplementary data are available at *Bioinformatics* online.

## 1 Introduction

Cross-linking technique in combination with mass spectrometry (MS) is commonly used to analyze protein structures and protein-protein interactions (Young *et al.*, 2000). Various computational tools have been developed to analyze cross-linking MS data. These tools include MS2Assign (Schilling *et al.*, 2003), xQuest/xProphet (Rinner *et al.*, 2008; Walzthoeni *et al.*, 2012), crux (McIlwain *et al.*, 2010), xComb (Panchaud *et al.*, 2010), Xlink-Identifier (Du *et al.*, 2011), Protein Prospector (Chu *et al.*, 2010; Trnka *et al.*, 2014), pLink (Yang *et al.*, 2012), MeroX (Götze *et al.*, 2014), MXDB (Wang *et al.*, 2014), Kojak (Hoopmann *et al.*, 2015), XlinkX (Liu *et al.*, 2015) and ECL (Yu *et al.*, 2016, 2017).

Compared with the identification of single peptides (Eng *et al.*, 1994; Perkins *et al.*, 1999; Tanner *et al.*, 2005), the identification of cross-linked peptides requires a much larger search space because we need to examine all pairs of candidate peptides. To be more precise, the search space of identifying cross-linked peptides is quadratic with respect to the number of candidate peptides in a database (Liu *et al.*, 2015). Therefore, it is computationally prohibitive to examine all possible candidate pairs in a large database.

To tackle this problem, MS-cleavable cross-linkers, such as disuccinimidyl sulfoxide (DSSO) (Kao *et al.*, 2011), BuUrBu (Müller *et al.*, 2010) and cyanurbiotindipropionylsuccinimide (CBDPS) (Petrotchenko *et al.*, 2011), have been introduced. With MS-cleavable cross-linkers, the identification of cross-linked peptides can be finished in linear time (Liu *et al.*, 2015). However, the time complexity is still quadratic when non-cleavable cross-linkers are used.

To reduce the quadratic time complexity when non-cleavable cross-linkers are used, most of the existing tools use some heuristic screening strategies to reduce the number of candidates. For example, pLink (Yang *et al.*, 2012) first conducts a coarse-grained scoring on all single peptides, and then selects top 500 of them as candidates for further fine-grained scoring. In the fine-grained scoring, only these 500 candidate peptides are paired together as candidates of cross-linked peptides. Kojak (Hoopmann *et al.*, 2015) uses a similar strategy to select only the top 250 of single peptides. Both tools limit the number of candidates when enumerating combinations, so the running time can be reduced to an acceptable level.

Instead of reducing the number of candidates, Chen *et al.*, 2001 proposed a general algorithm for identifying cross-linked peptides and provided theoretical analysis of their method. Their method can be decomposed into two stages. In the first stage, they generate possible solutions that satisfy the requirement of precursor mass. In the second stage, they measure the similarity between the experimental mass spectrum and each pair of peptides. Because of the large number of combinations, the authors suggested a speed-up implementation by removing low-score candidates in measuring similarities of single peptides. This is also a screening strategy.

Yu *et al.*, 2017 proposed an algorithm whose time complexity is linear with respect to the number of peptides in a database to solve the problem described above. They use the additive property of the scoring function, and a binning strategy to assign peptides into small bins according to their mass. Afterwards, they search and record the peptide with the maximum score in each bin, and enumerate bin pairs to figure out the most similar cross-linked peptides. As the number of bins can be defined beforehand, the computational cost on enumerating bin pairs will be fixed.

However, their method suffers from the following issues. The time complexity of enumerating bins in ECL2 is 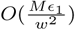, where *M* is the total range of peptide mass, *ϵ*_1_ is the MS 1 tolerance and *w* is the MS 1 bin width (Yu *et al.*, 2017). This time complexity can be rewritten as 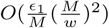, where 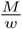 is the total number of bins, and 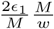 is the number of candidate bins within the MS1 tolerance range. Both are proportional to 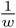, so the time complexity of their method is still quadratic with respect to the MS 1 mass precision 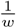. Since peptide combinations are replaced by bin combinations, i.e., the time complexity 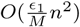 by enumerating peptide pairs (*n* peptides and average 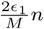 candidate peptides within the MS1 tolerance range) becomes 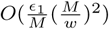 in ECL2, the time complexity is still quadratic with respect to the objects being enumerated. That is to say, the high complexity issue of enumerating combinations has not been fully solved. Besides, because the binning strategy is used in ECL2, the precursor mass constraint may not be strictly satisfied.

In this work, we propose a new linear-time algorithm to solve the problem of identifying cross-linked peptides. The proposed algorithm not only overcomes the above issues, but also shows an novel advantage on reducing the computational cost on scoring peptides. By using the additive property of the scoring function and the data structure of double-ended queue, the proposed algorithm exhaustively searches cross-linked peptides in linear time. The time complexity of the proposed algorithm is not only linear with respect to the number of peptides in a database, but also constant with respect to the MS 1 tolerance. We implement this algorithm in a tool named Xolik. The correctness proof and the time complexity analysis are given in the Supplementary Notes. Experiments using synthetic and empirical datasets show that Xolik outperforms existing tools in terms of running time and statistical power.

## 2 Methods

### 2.1 Problem formulation

Given a scoring function that measures the similarity between the query MS2 spectrum and the candidate cross-linked peptides, we have the following cross-linking problem:

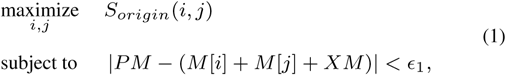

where *PM* is the precursor mass, *XM* is the mass of the cross linker, *ϵ*_1_ is the MS 1 tolerance, and *i* and *j* are peptide indices. For the *i*th peptide, *M*[*i*] is its mass. *S_origin_* (*i, j*) is the similarity score between the query MS2 spectrum and the cross-linked peptides formed by the *i*th and *j*th peptides.

Suppose the scoring function satisfies the following additive property:

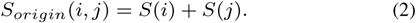

*S*(*i*) and *S*(*j*) denote the scores of the *i*th and *j*th peptide, respectively, with a pseudo-modification at the cross-linked site (Yang *et al.*, 2012). This additive property means that we are able to decouple the score of the cross-linked peptide as a sum of two scores corresponding to two single peptides. Therefore, the cross-linking problem becomes

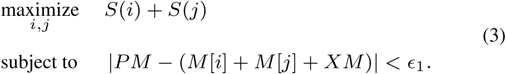

With the additive scoring function, the proposed algorithm finds the cross-linked peptides in linear time with respect to the number of candidate peptides in the database.

### 2.2 Linear-time search algorithm

Any scoring functions satisfying the above additive property can be used in our algorithm. In the default implementation, we use a modified version of the XCorr scoring funtion (Eng *et al.*, 2008). The modification is that we don’t incorporate theoretical ions from two peptides if they are within the same MS2 bin.

To identify cross-linked peptides from MS data, peptides in a sequence database are first digested *in silico* and sorted based on the mass in advance. (Sorting the mass requires *O*(*n* log *n*) time. However, since it is done offline, we don’t include it in the time complexity of analyzing a spectrum.) With the proposed algorithm, finding the cross-linked peptides given a query MS2 spectrum takes *O*(*n*) time. We only calculate the similarity score between the query spectrum and a candidate if the comparison between them is necessary. This allows us to reduce the computational cost on scoring because we merely score peptides that are necessary to solve the problem, rather than blindly score all candidates in the whole database. In the next subsetction, we will describe the benefits of this strategy. To search the peptide pairs that satisfy the precursor mass constraint, we use three pointers (i.e., *I_f_, I_bf_, I_be_*) to denote the current examined peptide, the lower bound and the upper bound of the index range of candidate peptides that satisfy the requirement of precursor mass, respectively (see Fig. 1 and Fig. 2). *I_f_* is initially pointed to the peptide with the smallest mass, *I_bf_, I_be_* are initially pointed to the peptide with the largest mass. We use a double-ended queue to maintain the order of scores compared in previous iterations. In each iteration, we examine one peptide and compute the range of the other peptide that satisfies the mass constraint. For each examined score, finding the other peptide with the maximum score in the valid range can be solved in a constant amortized time with the help of the double-ended queue. Therefore, after all iterations, the maximum score among all peptide pairs corresponding to the query MS2 spectrum is available in linear time with respect to the number of candidate peptides in the database.

**Fig. 1:**
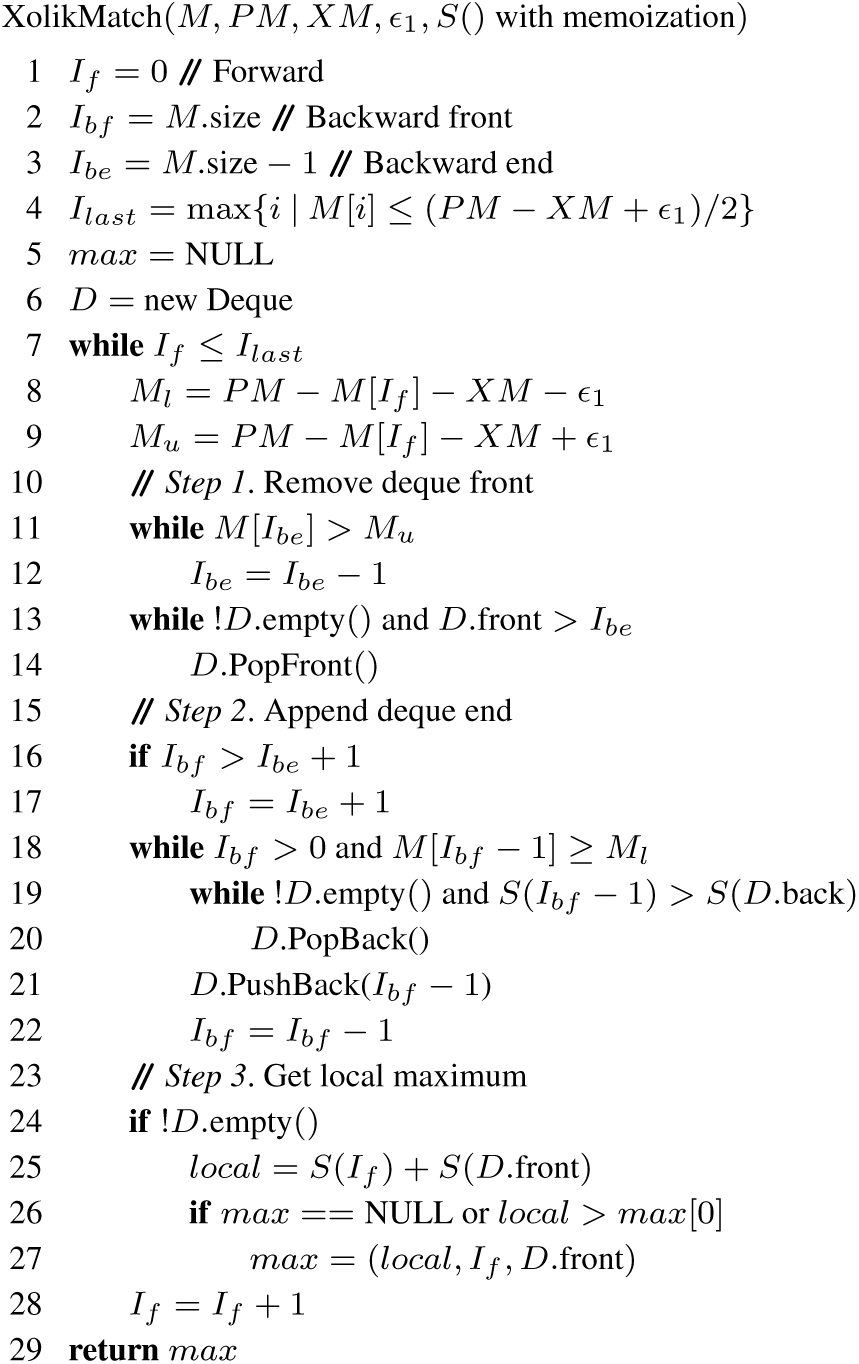
The pseudocode of the algorithm that finds the cross-linked peptides in linear time. Given the mass of all candidate peptides *M* (*M* is sorted in ascending order beforehand), the precursor mass *PM* of the query MS2 spectrum, the cross-linker mass *XM*, the MS1 tolerance *ε*_1_, and the scoring function *S*() with memoization, the XolikMatch algorithm finds the cross-linked peptides in linear time. The basic idea of this algorithm is to use a double-ended queue *D* to store the order of scores compared in previous iterations.

**Fig. 2:**
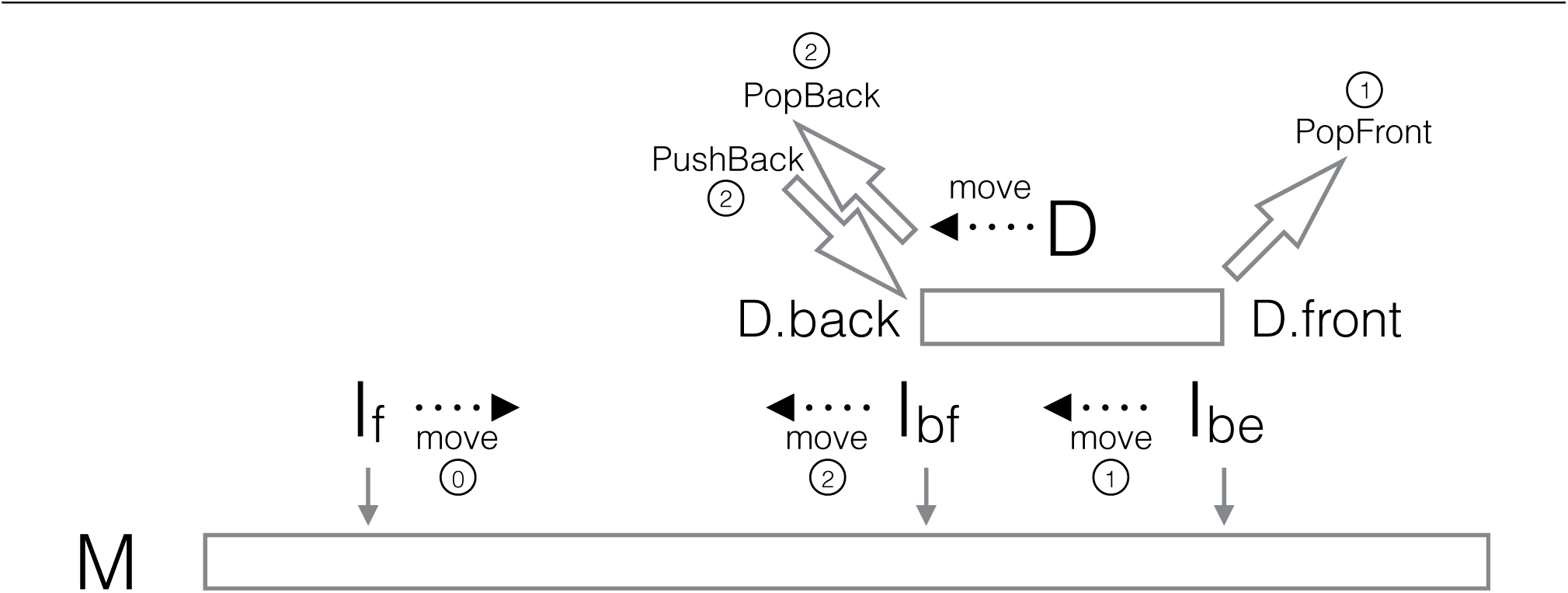
Graphical illustration of the linear-time algorithm in each iteration. The graph shows the change of program states starting from the previous iteration at Line 24 of the pseudocode (Fig. 1). *I_f_* indicates the position of the current examined peptide. *I_bf_* and *I_be_* indicate minimum and maximum indices that meet the requirement of precursor mass, respectively. *D* is a double-ended queue (deque), in the opposite direction to *M* (the sorted mass array). The circled numbers denote the steps. In Step 0, *I_f_* moves right (the end of the previous iteration). Step 1 adjusts *I_be_* and pops elements out from the front of the deque *D*. Step 2 moves *I_bf_* to the left and pushes elements into the back of the deque *D* if the elements are with scores less than or equal to the back of the deque *D*, or recursively pops elements out from the back of the deque *D* if the score of the new element is greater until the new element meets the requirement to be pushed into the back of the deque *D*. With the completion of Step 2, the front of the deque *D* is the index that indicates the maximum score in the range [*I_bf_, I_be_*].

The improvement of the proposed linear-time algorithm comparing with the algorithm proposed in ECL2 (Yu *et al.*, 2017) is at the overhead on enumerating bin pairs. The time complexity of ECL2 is 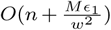, where *O*(*n*) is the time complexity of scoring peptides and assigning peptides into bins, and 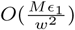 is the time complexity of enumerating bin pairs. Xolik directly solves the problem in *O*(*n*) time, without any binning strategies. Therefore, the overhead 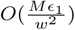 in ECL2 is suppressed in Xolik. Besides, because no binning strategy is used, Xolik doesn’t relax the constraint of the precursor mass.

### 2.3 Lazy evaluation on scoring single peptides

A nice property of the algorithm is that, when we retrieve a similarity score of a peptide, we are sure that this peptide must be in some valid solutions of cross-linked peptides that meet the requirement of precursor mass. So the score of this peptide is necessary for the algorithm to figure out the most similar cross-linked peptides. At the same time, if a peptide is not in any valid solutions of cross-linked peptides, we will never retrieve the score of it. Therefore, we can postpone the computations of similairty scores of single peptides until we retrieve the scores for comparison. This lazy evaluation strategy saves resources on computing the similarity scores of those “useless” peptides. Along with the memoization on the computed scores, all similarity scores will be computed at most once. To implement the memoization technique, we only need an additional cache layer to store the flags indicating whether scores have been calculated or not. This additional layer only requires a small amount of memory space. For a typical number of peptides in a large database, e.g., 2,000,000 peptides, a naive implementation using 64-bit int as flags merely needs 8 bytes × 2000000 ≈ 16 MB extra memory.

## 3 Results

### 3.1 Running time validation

In this subsection, we will examine whether Xolik can complete the identification task in linear time. As a comparison, we also run ECL (Yu *et al.*, 2016) and ECL2 (Yu *et al.*, 2017) to finish the same tasks. We use a MS data file (20111221_ananiav_DPDS_lib1_90min_CID35.mzXML) from a synthetic dataset (Wang *et al.*, 2014) to search against random databases with different sizes (i.e., different number of proteins). Random databases are generated by randomly selecting proteins from the human protein database, with protein numbers ranging from 100 to 20000. The MS1 tolerance is set at 50 ppm. Also, to validate the effect when the MS1 tolerance changes, we also search the data file against a random databases containing 10000 proteins, with MS1 tolerance ranging from 50 ppm to 500 ppm. The MS2 bin size for XCorr is set at 0.5 Da (roughly 0.25 Da MS2 tolerance), and the maximum number of missed cleavage is 2. The mass range of candidate peptides is [500 Da, 5000 Da]. In this experiment, we want to compare the running time in a common scenario, so we set BS3 as the cross-linker and enable the E-value estimation, even though the MS spectra in the data file are synthetically cross-linked by SS-bond (Wang *et al.*, 2014). We also search against the decoy database, which is constructed by reversing the protein sequences in the target database. The MS data file contains 16557 MS2 spectra. The running time versus the database size is shown in Fig. 3a, and the running time versus the MS1 tolerance is shown in Fig. 3b. All tools are deployed on an Intel Core i5 3.20GHz Linux desktop computer with 16GB memory, which is a regular PC with a basic configuration, and run in 4 threads (if applicable) with all memory assigned to the process.

**Fig. 3:**
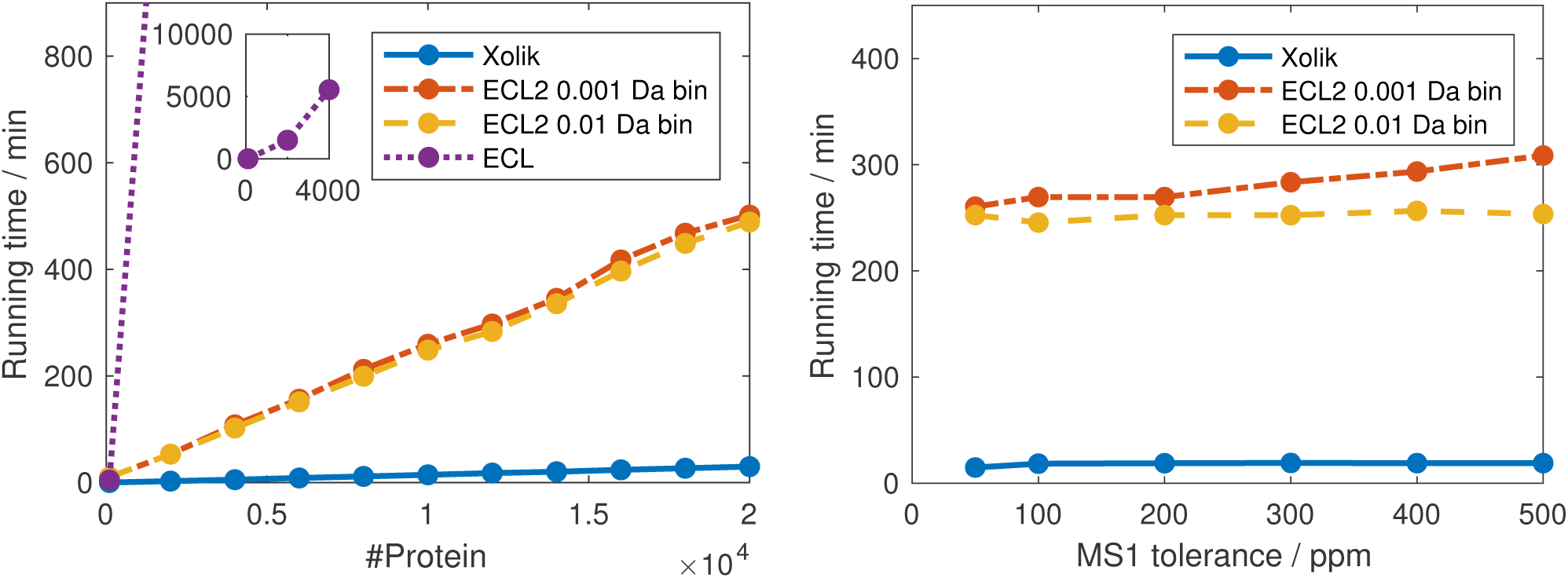
The running time v.s. the database size (#Protein) (Fig. 3a) and the MS1 tolerance (Fig. 3b). (a) The databases are searched with MS1 tolerance at 50 ppm and MS2 bin size for XCorr at 0.5 Da. As the number of proteins increases, the running time of Xolik increases linearly. When searching a database containing 20000 proteins, Xolik takes about 30 min. For ECL2, the curves in both settings increase linearly. However, the running time of ECL2 with a smaller MS1 bin width is less than that with a larger MS1 bin width. The curve of ECL increases quadratically, which is consistent with its time complexity. Not only because of its quadratic time complexity, but also because of the lack of multithreading, ECL spends around 5540 min (92 hours) searching a database with 4000 proteins. (b) The database contains 10000 random proteins. As MS1 tolerance increases, Xolik roughly maintains the same running time. For ECL2, as the MS1 tolerance increases, the running time with 0.001 Da MS1 bin width increases linearly. In contrast, the running time with 0.01 Da MS1 bin width only slightly increases, which reflects that enumerating 0.01 Da MS1 bins only costs one-hundredth comparing with enumerating 0.001 Da MS1 bins. ECL is skipped in this experiment because it cannot handle 10000 proteins within an acceptable period of time.

Fig. 3a shows that the running time of ECL increases quadratically, the running time of ECL2 increases linearly, and the running time of Xolik also increases linearly, with respect to the number of proteins in the database. All tools are consistent with their time complexities, respectively. Most notably, Xolik searches a MS data file against 20000 proteins in around 30 minutes on a regular PC. This allows us to search against a large database within an acceptable period of time, which especially benefits the analysis of complex protein samples. We also show the performance of Xolik in searching a real dataset against a large protein database in the last experiment. The running time of Xolik is stable when the MS1 tolerance *ϵ*_1_ increases. In contrast, the running time of ECL2 with 0.001 Da MS1 bin width increases roughly linearly. Comparing with the running time with 0.01 Da MS1 bins, when the MS1 tolerance is large, e.g., 500 ppm, one additional order of magnitude on the MS1 mass precision requires 55 min extra running time for ECL2.

We provide the theoretical analysis for Xolik’s linear-time algorithm in the Supplementary Notes. It proves that it is indeed a linear-time algorithm and the time complexity will not change with respect to the MS1 tolerance.

### 3.2 Analysis of synthetic disulfide-bridged peptides

We run Xolik on a synthetic disulfide-bridged peptide dataset (Wang *et al.*, 2014) to examine its statistical power. As a comparison, we also run Kojak, pLink and ECL2 on the same dataset. Since pLink reports few identifications (163 peptide-spectral matches (PSMs)) and ECL2 currently does not support the customization on the cross-linked site (SS-bond links at Cysteine (C)), we skip the comparison with pLink and ECL2. The dataset we analyze contains three synthetic peptide libraries, which have the following three specific sequence patterns:

- K[AW][DE]F[VSHY]A[DY]SCVA[KR]
- [TW]A[LE]H[FV]SCVT[PSGY]F[KR]
- [WA]VK[FL]C[DE]T[VSGY]FA[KR]

Please refer to the original paper (Wang *et al.*, 2014) for a detailed description of the sample preparation and the XL-MS analysis.

To validate the statistical power of Xolik, we search these libraries against a target database containing all peptide sequences matching the above patterns. Total 704 peptide sequences are in the target database. The MS1 tolerance is set at 50 ppm and the MS2 bin size for XCorr is set at 0.5 Da. The maximum number of missed cleavage is 2. The mass range of candidate peptides is [500 Da, 5000 Da]. The cross-linker used in the database search is SS-bond. The decoy database is constructed by reversing the protein sequences in the target database, and the false discovery rate (FDR) is controlled at 5%. The FDR controlling procedure implemented in Xolik is the same as the method used in xQuest/xProphet (Walzthoeni *et al.*, 2012) and pLink (Yang *et al.*, 2012). For Kojak, we follow the whole identification procedure using Percolator (Kall *et al.*, 2007) to control the FDR of the findings. Since Kojak will output multiple PSMs for one spectrum, for a fair comparison, we only keep the top match for each query spectrum on the controlled identifications. Because the sequences in the database follow certain patterns, the basic assumption of the randomness for E-value estimation is violated. Controlling FDR on E-value significantly degrades the performance, even though most of the dropped records can be validated by the above sequence patterns (data not shown). Therefore, we run Xolik without E-value estimation in this experiment. Both tools run in an Intel Core i5 3.30GHz Windows desktop computer with 12G memory in 4 threads. The detailed comparison on the search results is shown in Table 1.

**Table 1.**
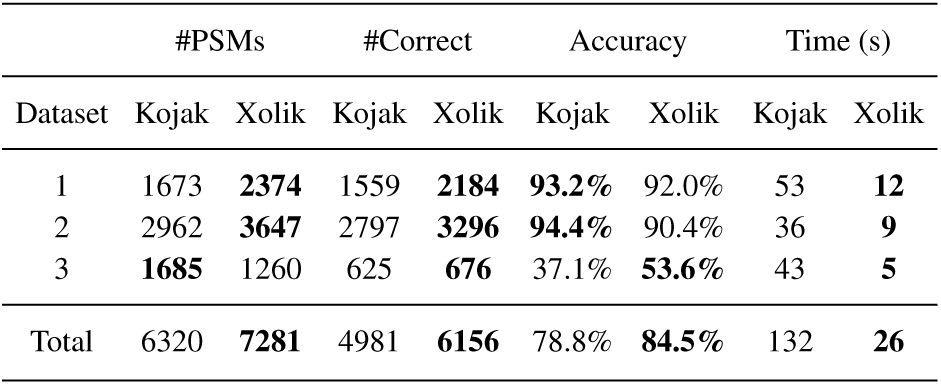
Comparisons between Xolik and Kojak on a synthetic disulfide-bridged peptides dataset (Wang et al., 2014). Both tools run on an Intel Core i5 3.30GHz Windows desktop computer with 12GB memory in 4 threads. Accuracy is defined as #Correct / #PSMs.

Each library only contains peptides following one specific sequence pattern, so we are able to evaluate the reported PSMs by comparing the identified sequences with the sequence pattern corresponding to the library. Only when both sequences of a cross-linked PSM match the sequence pattern is the reported peptide pair considered as a correct identification.

As shown in Table 1, Xolik identifies more PSMs than Kojak in total. After manual examination on the identified sequence patterns, we find that Xolik also identifies more correct PSMs than Kojak in all datasets. This indicates that Xolik has a higher statistical power. Even though Xolik has lower accuracy than Kojak in the first two datasets, Xolik is more accurate than Kojak on average among all datasets. Moreover, Xolik searches all candidate peptides exhaustively but still runs faster than Kojak.

### 3.3 Analysis of *Escherichia coli* 30S and 50S ribosomal subunits

To illustrate the performance on real datasets, we run Xolik on a *E.coli* ribosome dataset (Lauber and Reilly, 2011). There are 48 mass spectra files analyzed in total. As a comparison, we also run ECL2, pLink and Kojak on the same dataset. All proteins in *E. coli* 30S and 50S ribosomal subunits are included in the target database (55 proteins), and the decoy database is constructed by reversing the protein sequences in the target database. The MS1 tolerance is set at 5 ppm, and the MS2 bin size for XCorr is set at 0.02 Da. The maximum number of missed cleavage is 2. We set a fixed modification +57.02146 at Cysteine (C), and no variable modification. The allowed mass range of candidate peptides is [500 Da, 5000 Da]. Since pLink cannot set the MS2 tolerance and the mass range of candidate peptides, we use the default setting when running pLink. The cross-linker used in the database search is diethyl suberthioimidate (DEST) (Lauber and Reilly, 2011). The E-value estimation and the multithreading are enabled if applicable. All tools are deployed on an Intel Core i5 3.30GHz Windows desktop computer with 12G memory. When running ECL2, all memory (12G) are assigned to the process. We control the FDR on the reported identifications at 5% using the methods bundled in each tool. The search results of all tools are shown in Fig. 4. As shown in the figure, Xolik shows significant improvement in terms of running time. Also, in terms of the number of identified PSMs, Xolik reports more PSMs than pLink and Kojak, and slightly outperforms ECL2. Although Xolik reports more PSMs than ECL2 in this experiment, it is not clear whether a strict precursor mass constraint will lead to an increment of identifications or not.

**Fig. 4:**
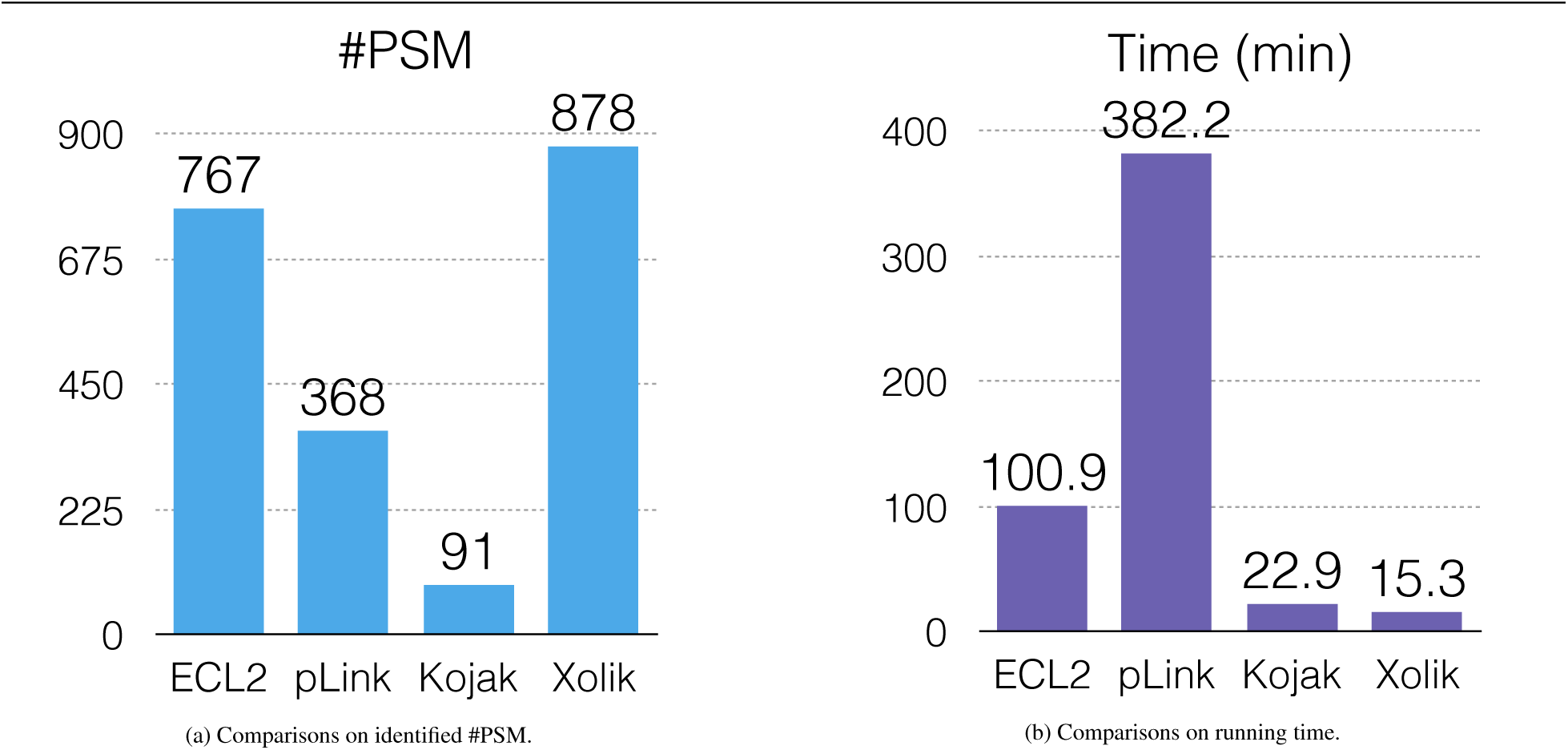
Identification results on the *E. coli* ribosome dataset (Lauber and Reilly, 2011) under 5% FDR control. All tools run on an Intel Core i5 3.30GHz Windows desktop computer with 12GB memory. Xolik reports more PSMs than other tools. Also, Xolik outperforms other tools in terms of running time. The difference between ECL2 and Xolik is only at the matching algorithm for pairing two single peptides. However, because the algorithm used in ECL2 relaxes the constraints of the MS1 tolerance, peptides outside the MS1 tolerance range are possibly reported by ECL2. Therefore, around the boundary of the tolerance range, ECL2 and Xolik may assign different labels to the query spectrum. This also affects the threshold determined by the FDR controlling algorithm. As a consequence, after controlling the FDR, the identification results between Xolik and ECL2 are partly different.

### 3.4 Analysis of *Homo sapiens* HeLa cell dataset

We also run Xolik on a human sample dataset (Makowski *et al.*, 2016) to evaluate the performance when searching a large protein database. This dataset contains around 3 × 10^5^ MS2 spectra, and the whole human protein database (downloaded from UniProt at 2016.03.10, total 20198 proteins) is used in the database search. The decoy database is constructed by reversing the protein sequences in the target database, and the FDR is controlled at 5%. The MS1 tolerance is set at 5 ppm, and the MS2 bin size for XCorris set at 0.02 Da. The maximum number of missed cleavage is 2. We set a fixed modification +57.02146 at Cysteine (C), and no variable modification. The mass range of candidate peptides is [500 Da, 5000 Da]. The cross-linker used in the database search is BS3. We also run ECL2, pLink and Koj ak on the same dataset for comparison. ECL2 cannot handle a database with more than 20000 proteins because the Java platform used by ECL2 requires more than 32G memory during the analysis. As a consequence, ECL2 spends most of time waiting for spare memory. Neither ECL2 nor pLink can finish the analysis within a week. Therefore, we only show the results of Xolik and Kojak. We enable the E-value estimation in Xolik and enable multithreading in both tools (8 threads). All tools are deployed on an Intel Core i7 3.50GHz Windows desktop computer with 32G memory. The result is shown in Fig 5. Xolik runs much faster than Kojak even though Xolik searches exhaustively. Also, Xolik reports more PSMs than Kojak.

**Fig. 5:**
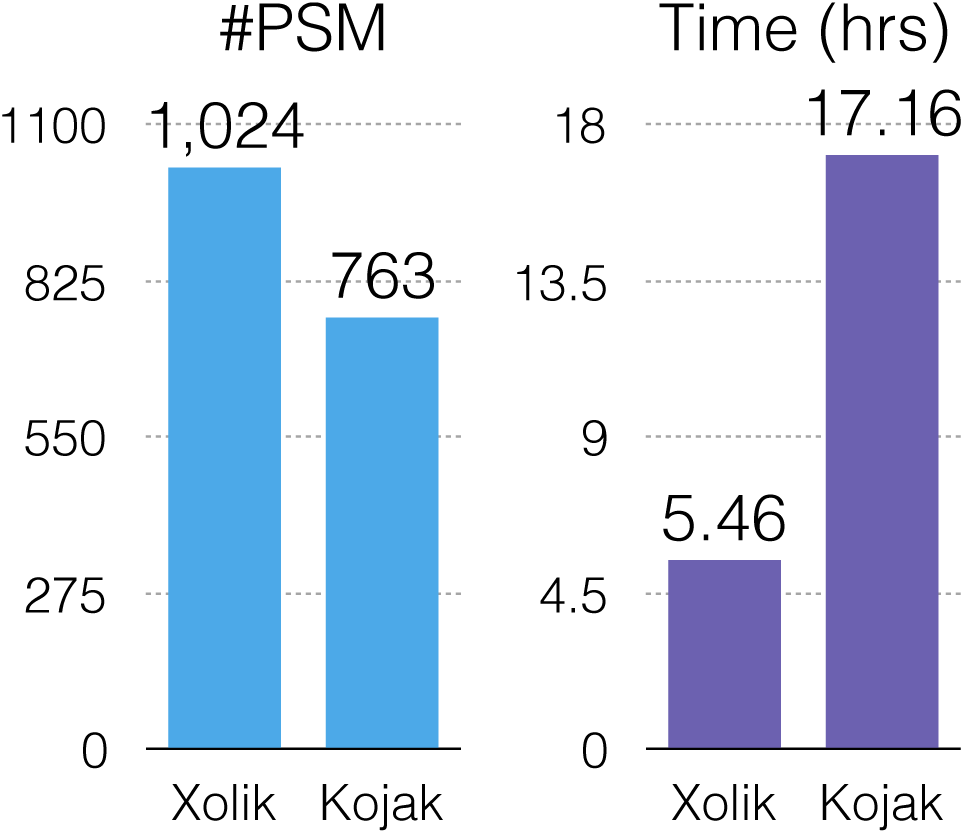
Identification results on the human protein dataset (Makowski *et al.*, 2016) under 5% FDR control. Both tools run on an Intel Core i7 3.50GHz Windows desktop computer with 32GB memory in 8 threads. Xolik outperforms Kojak in terms of running time, even though Xolik searches cross-linked peptides exhaustively. In terms of identified PSMs, Xolik also outperforms Kojak by reporting more PSMs at the same FDR level.

## 4 Conclusion

In this paper, we propose a linear-time algorithm for finding cross-linked peptides with maximum paired scores in a protein sequence database. It is implemented in a tool named Xolik. Instead of adopting screening strategies to reduce the search space, the proposed algorithm exhaustively searches the original search space by utilizing the additive property of the scoring function. Using the double-ended queue to store the order of scores compared in previous iteration, the proposed algorithm achieves the linear time complexity with respect to the number of candidate peptides in the database. Moreover, utilizing the lazy evaluation strategy together with the memoization technique, Xolik further reduces the computational cost on computing similarity scores. Experiments on a synthetic dataset and two empirical datasets show that Xolik outperforms existing tools in terms of running time and statistical power.

